# Insight into the mechanism of protein thermostability based on the residue interaction degrees

**DOI:** 10.1101/135319

**Authors:** Huihua Ge, Yunmeng Chu, Guangya Zhang

## Abstract

Understanding the basis of protein thermostability raises a general question: which residue with specific interaction degrees is more important to the protein thermostability? A strictly selected dataset of 131 pairs of thermophilic (TPs) and mesophilic proteins (MPs) was constructed. There were 6.4% and 8.4% of the total residues in sequences did not interact with others in TPs and MPs. The amino acid contents in sequences are closest to those with the interaction degrees of 3 according to the Chi-squared distances. Only Glu, Gln and the amide residues showed significant differences in sequences, which was the same as identified at low residue interaction degrees. However, we observed significant Phe, Lys, Leu, Gln and the charged, aliphatic, aromatic, positive charged and small residues at high interaction degree. Among them, Phe was rarely reported previously although aromatic residues were well-known contributor to protein thermostability. Finally, we took aspartate transcarbamylases as an example to explain how a residue with various interaction degrees contributed differently to their thermostability. Our results clearly demonstrated the differences of amino acids in sequence between TPs and MPs could only represent those involved in low interaction degrees. Much more residues with significant differences existed at high interaction degrees even if they had few significant amino acids in sequences. The interaction degree-based method should be an alternative tool in extracting valuable eigenvalues for predicting proteins attributes in bioinformatics. It could also provide a new perspective for studying the thermostability of proteins and engineering novel thermostable proteins.

**List of abbreviations:**

TPs: thermophilic proteins
MPs: mesophilic proteins
OGT: optimal growth temperature
ASA: absolute surface areas

## Introduction

Elucidating the mechanisms of the stability of thermophilic proteins is an important field in computational biology. Researcher have found many factors contributed to the thermostability of proteins, though some of them are still debated [1-5]. However, it is widely accepted amino acids play an important role in thermostabilization. Researchers have observed significant differences of amino acids between TPs and MPs in the proteome [6], the surface and interior of the proteins [7] and various secondary structures [8]. Besides, they adopted many classifiers to discriminate thermophilic and mesophilic proteins based on the amino acid composition [9-11]. The high overall prediction accuracy proved the amino acid composition was an effective eigenvalue to distinguish these two types of proteins.

However, based on the known facts, it is difficult to answer the following questions: does the identical residue with different interaction degrees contribute equally to the protein thermostability? If not, which residue with specific interaction degrees is more important to the thermostability of proteins? Can we identify those amino acids showing significant differences according to their interaction degrees? Despite many works devoted to the search for amino acid differences between TPs and MPs, these questions were not mentioned and solved. At least, the works existed currently did not provide accurate fingerprints about the interaction degrees of amino acids when compared their features between TPs and MPs.

From a theoretical perspective, the 3D structure of a protein was determined by a delicate balance of different interactions between residues. Without a doubt, the amino acids contribute to the stability of a protein *via* their mutual interactions. As we all know, many interactions such as the hydrogen bonds, salt bridges, π-cationic interactions and so on play important roles in the thermostability of proteins [12, 13]. Sometimes, an amino acid may involve in more than one kinds of interactions in a protein, and sometimes it may not interact with others in the same protein. For example, thermophilic proteins showed higher frequencies of arginines in sequence or in exposed state, they contributed positively to the thermostability [14, 15]. However, there are many arginines in protein sequence, they have different interaction with other residues depending on their positions and those residues involved. For instances, there are two arginines which do not interact with others (e.g. Arg-12 and Arg-152 in 1ML4, 1ML4 is the aspartate transcarbamylase from the hyperthermophilic archaeon *Pyrococcus abyssi* [16].), while three arginines have nine interactions with others (e.g. Arg-107, Arg-168 and Arg-297 in 1ML4). Do they contribute equally to the protein thermostability? If not, how do we differentiate their role to the thermostability of proteins? Unfortunately, few works have addressed it although researcher have studied protein thermostability for a long time. From an applicative perspective, the amino acids with specific interaction degrees that show significant differences between TPs and MPs are quite definite, especially those with high interaction degrees. Thus, it is convenient to target these residues when engineering for thermophilic proteins. Meanwhile, the outcomes will be improved and more trustable.

Here, we strictly selected thermophilic and mesophilic homologs and created a dataset containing 131 pairs of proteins. They had high sequence identities and only few residues showed significant differences in sequences. Our first purpose is to identify more amino acids at some interaction degrees that contributed significantly to the thermostability of proteins. The second purpose is to differentiate the role of the same amino acid with various interaction degrees to the protein thermostability.

## Materials and methods

### Dataset construction

The original dataset of 252 pairs of TPs and MPs was compiled by Taylor *et al*. [17]. However, many identical proteins have two or more different mesophilic or thermophilic homologues in this dataset. To reduce the computational bias, we kept only one pair of them each. The criterion is a). The homologues have the higher sequence identity; b). Their source organisms have higher optimal growth temperature (for TPs) or have lower optimal growth temperature (for MPs); c). The sequence length is closer; d). Discarding the sequences with less than 100 amino acids. We got 112 pairs of homologues from this dataset. Then we gather another 19 pairs of homologues, which was completely different. Finally, we got 131 pairs of proteins. The information containing the PDB IDs, the length, the resolution of the structure, the source organisms, their corresponding optimal growth temperature (OGT), and the sequence identity were listed in the supplementary table S1. There were 40384 and 41609 amino acids in the sequences of TPs and MPs. Meanwhile, there were 2578 and 3481 amino acids did not interact with others, accounting 6.4% and 8.4% for TPs and MPs.

### Interaction degree

The interactions calculated here include hydrogen bond, Van der Waals, disulphide bond, salt bridge, π-π stacking and π-cation according to the Residue Interaction Network Generator (RING) at the RING 2.0 web server [18]. The cutoff distances for hydrogen bond, Van der Waals, disulphide bond, salt bridge, π-π stacking and π-cation were 3.5 Å, 0.5 Å, 2.5 Å, 4.0 Å, 6.5 Å and 5.0 Å, respectively [18]. The interaction degree (*D*) of an amino acid means the number of interactions it interacts with others. We divided the degrees into three groups, namely, low degree (1 ≤ D ≤ 4), medium degree (5 ≤ *D* ≤ 8) and high degree (*D* ≥ 9). Their proportion was 59.1%, 32.4% and 2.1% for TPs, while 59.4%, 30.3% and 1.9% for MPs, respectively.

### Definition of the Chi-squared distances

We might count the frequency of each residue at a specific interaction degree and compared them with what we would expect (the frequency of each residue in the sequence) by means of Chi-squared distances. If there were no preference of the residues at a specific interaction degree, the frequencies of the residues at the specific interaction degree (observed) would be the same as those in the sequence (expected). Thus, the distances would be zero. Therefore, the Chi-squared distances can help us make decisions about whether the observed outcome differs significantly from the expected outcome.

The Chi-squared distances between sequence and different interaction degrees was defined as following according to Abdullah’s work [19]:

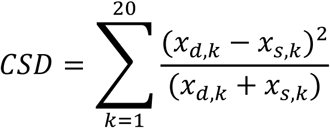

Where *k* is the 20 amino acids in proteins, *x*_*d*,*k*_ is the average amino acid composition at a specific interaction degree, *x*_*s*,*k*_ is the average amino acid composition in sequences.

The critical value for the chi-square in this case is 30.144 (with the degree of freedom of 19). If the calculated Chi-square value is equal to or greater than this critical value, we can conclude that the probability of the null hypothesis being correct is 0.05 or less, which is a very small probability indeed! Then we reject the null hypothesis. For example, our calculated value of 79.511(the distances between sequence and the interaction degree of 10 for TPs, as shown in figure 2.) is greater than the critical value of 30.144. We therefore reject the null hypothesis, and conclude that there is a significant difference in amino acid composition between the sequence and residues at the interaction degree of 10 in TPs.

Similarly, for the Chi-squared distances between sequence and different secondary structure, the distances were defined as:

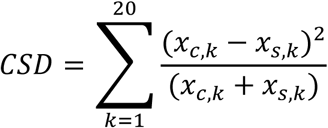

Where *k* is the 20 amino acids in proteins *x*_*c,k*_ is the average amino acid composition at a specific secondary structure (α-helix, β-sheet and coil), *x*_*s,k*_ is the average amino acid composition in sequences.

For the Chi-squared distances between the different relative solvent accessibility and the sequences, the distances were defined as:

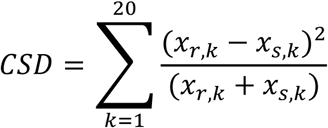

Where *k* is the 20 amino acids in proteins, *x*_*r,k*_ is the average amino acid composition at a specific relative solvent accessibility state. *x*_*s,k*_ is the average amino acid composition in sequences. According to relative solvent accessibility, the states of amino acids in proteins were classified as being in one of the four classes: completely buried (0-4% exposed), partly buried (4-25% exposed), partly exposed (25-50% exposed) and completely exposed (50+% exposed) [20].

### Significant amino acids (or amino acid groups) detection

The differences of amino acid (or amino acid groups) composition between TPs and MPs were defined as:

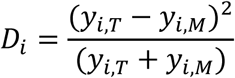

Where, *y*_*i,T*_ and *y*_*i,M*_ are the amino acid (or amino acid groups) composition of TPs and MPs, *i* means the specific degree (or degree range) or structure (e.g. the secondary structure such as helix, sheet or coil).

If *D*_*i*_ > 0.2, *y*_*i*_ is identified as significant amino acid (or amino acid group). We classified the amino acid as the following groups according to their physicochemical properties [21, 22]. The amino acid groups include the charged, aliphatic, aromatic, polar, neutral, and hydrophobic, positive charged, negative charged, tiny, small, large, sulfur and the amide.

## Results and Discussion

### The Chi-squared distances between the sequence and interaction degrees based on amino acid composition

To see if the Chi-squared distances can reflect the amino acids preferences from sequences, we calculated the Chi-squared distances between sequence and various secondary structure or relative solvent accessibility of TPs and MPs and showed the results in figure 1. We observed the Chi-squared distances of various secondary structure and the partial exposed and buried were between 2.5 and 12, while the distances of the exposed and buried were between 20 and 24. Among them, the exposed state shows the most distinct differences between TPs and MPs. This means the amino acid contents in the exposed part (protein surface) is the most biased when compared with those in the sequences. The Chi-squared distances were 23.55 and 21.38 for TPs and MPs. The absolute difference value was 2.17, which showed the most obvious differences between TPs and MPs. Researchers have found the most of the amino acid composition bias between TPs and MPs came from the protein surface (or the exposed part) [4, 7, 23]. Our results confirm it by the precise data of Chi-squared distances. As for the secondary structure, no significant differences observed between TPs and MPs in terms of Chi-squared distances, which is also in accordance with previous result [8, 24]. The amino acids in the β-sheet shows the maximum deviation over the sequence, while α-helix the minimum. The Chi-squared distances of β-sheet is 9.17 and 8.4 in TPs and MPs, while 4.34 and 4.15 for α-helix, respectively. However, the distances difference of the coil is more obvious between TPs and MPs. The distances are 6.33 and 5.36, with an absolute difference value of 0.98.

**Fig. 1.**
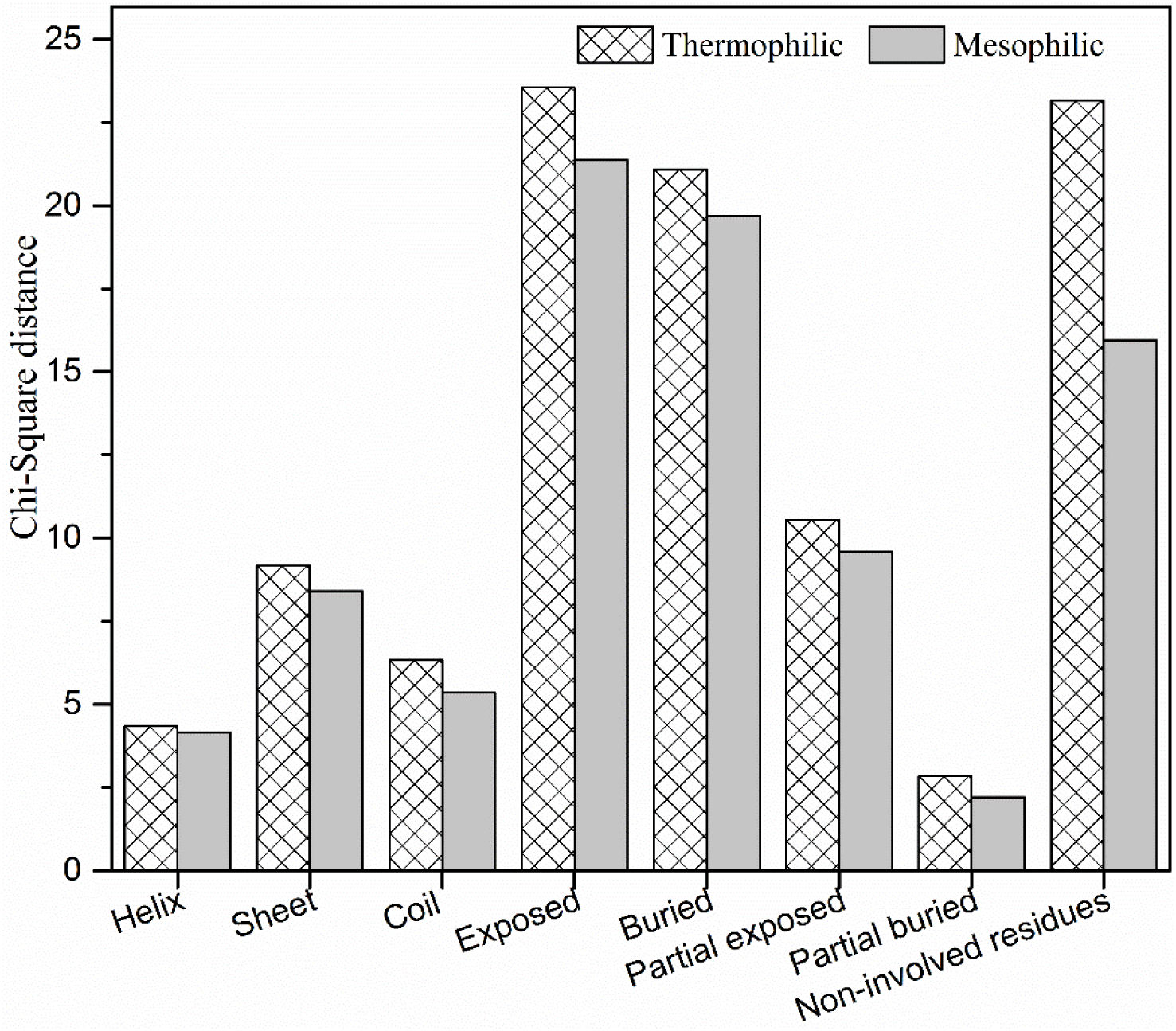
The Chi-squared distances between sequence and secondary structure, relative solvent accessibility and non-involved amino acids of TPs and MPs.

As the Chi-squared distances is the summation value of the distances for each of the 20 amino acids, it can also identify the amino acid that contributed the greatest to the distances. The residue contributed the greatest to the distances is the most biased one when compared with its content in sequence. For instances, the primary contributor to the distances between the protein surface and sequence is Leu in both TPs and MPs (accounting 16.9% and 16.6%), as the content of Leu is much less in the exposed part than in the sequences. The primary contributor for the distances between the buried part and sequence is Lys in both TPs and MPs (accounting 19.2% and 20.1%); as the content of Lys is much less in the buried part. As we all know, Leu is a hydrophobic residue, which involves in the hydrophobic interactions in the buried part of a protein; meanwhile, Lys is a polar charged residue, which should be abundant in the exposed part of a protein [4]. Thus, they become the most biased residues in the exposed and buried parts. For the secondary structure, Gly is the primary contributor to the distances between the α-helix and sequence, and Val the primary contributor between the β-sheet and sequence. According to Chou & Fasman’s secondary structure parameters, Gly is the strongest α-helix breaker, while Val the strongest β-sheet admirer [25]. Therefore, much less Gly in α-helix and more Val in β-sheet was observed and they became the primary contributor to the Chi-squared distances. Based on these results, we may conclude the Chi-squared distance we proposed here is biologically meaningful and convincing. Next, we will analyze the Chi-squared distances between the sequence and various interaction degrees.

We computed the average amino acid composition of the sequence and various interaction degrees and calculated the Chi-squared distances between them as showed in figure 2. We can see both TPs and MPs have very similar trends. The distances decreased before degree 3 and increased gradually after degree 4. The distances were the minimal when the degree is 3 and 4. They were 2.12 and 2.49 for TPs, while 2.00 and 2.16 for MPs, respectively. This means the amino acid composition at degree 3 and 4 approximates that of the sequence. The most important contributor to the distances comes from Phe (account 34.0% and 26.0% for TPs and MPs) at degree 3 and Gly (account 52.2% and 49.1% for TPs and MPs) at degree 4. Besides, the distance values of TPs were smaller than those of MPs when the degree is less than or equal to 6, and they became larger when the degree is greater or equal to 7. This means the usage of amino acids at relatively high degree of MPs biased more from their sequences. When we sorted the interaction degrees into low, medium and high, the Chi-squared distances of TPs and MPs were 66.49 *vs* 75.43 at high interaction degree, which was greater than the critical values of 30.144. This means there is a significant difference in amino acid composition between the sequence and residues at the high interaction degree. However, the Chi-squared distances at low (3.88 *vs* 3.40) and medium degree (17.77 *vs* 17.98) were less than the critical values, which means no significant differences observed. The most important contributors to the distances were Phe (account 18.0% and 21.2% for TPs and MPs) at high degree, while Gly (account 33.9% and 30.6%) at medium degree and Phe (account 19.8% and 21.2%) at low degree, respectively. Interestingly, the content of Phe in sequence is less than that of high degree, while more than low degree. As we know, Phe has a benzene-like ring structure in its side chain and it is a highly hydrophobic amino acid. It could involve in many kinds of interactions for maintaining conformational stability in protein structure [26]. Consequently, the frequency of Phe in high interaction degrees is much higher than that in sequence.

**Fig. 2.**
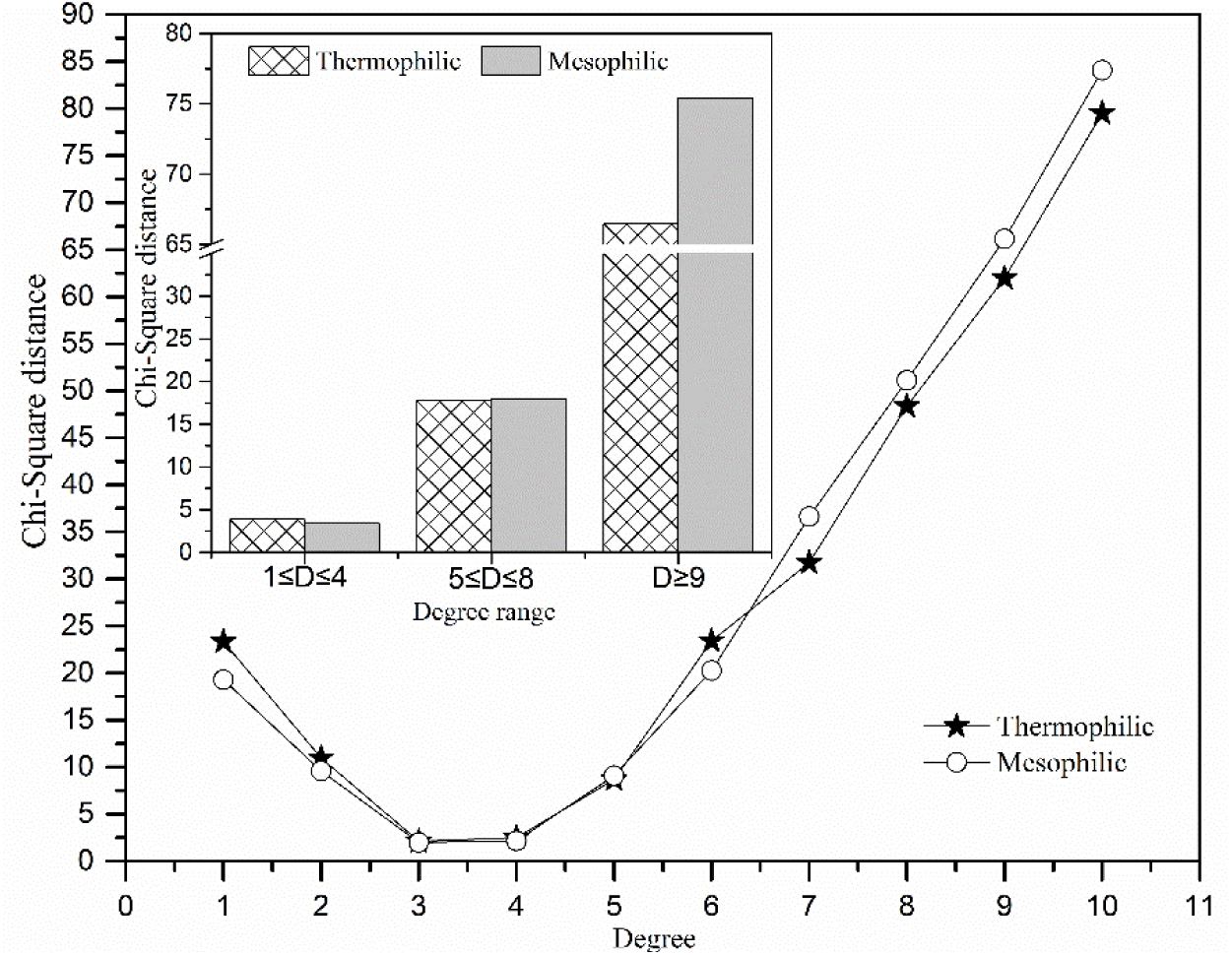
The Chi-squared distances between sequence and different residue interaction degrees.

If a residue in the sequence did not interact with others, they should scarcely have contributions to the thermostability of proteins. We identified those residues who did not involve in the interaction and calculated the contents of them in TPs and MPs. We found the content was 6.4% and 8.4% for TPs and MPs, respectively. The content is 2% less in TPs, meaning more amino acids in TPs involve in the interactions with other residues. The Chi-squared distances of TPs and MPs between sequence and the non-involved residues (amino acids do not involve in the interactions) were shown in figure

1. The distances for TPs and MPs were 23.16 and 15.96, respectively, which is also less than the critical value.

When compared the Chi-squared distances of the high interaction degree with those of the exposed or buried part (as shown in figure 1 and 2), we found the distance of the high interaction degree was much more. This means the usage of the amino acids at high interaction degree in more biased than that on the exposed and buried part of TPs and MPs. As mentioned above, the exposed and buried part were distinctively differ between TPs and MPs, which has been proved by many earlier researchers [1, 2, 4, 5, 7]. Here, we found the amino acids were far more biased at the high interaction degrees, we believe the differences between TPs and MPs are extremely significant at high interaction degree when compared with the surface and core part of proteins. Next, we will discuss the amino acid differences between the TPs and MPs according to their interaction degrees in detail.

### The amino acids showing significant differences between TPs and MPs at various interaction degrees

To prove our definition about the significant amino acids was effective, we firstly listed the amino acids showed significant differences between TPs and MPs in sequences and various relative solvent accessibility and secondary structure in table 1. We found there were significantly more Glu while fewer Gln and amide residues in the sequences of TPs. This is consistent with previous results [15, 27]. As for the relative solvent accessibility, we found the most obvious differences were in the exposed parts. There were significant more Glu, Arg, charged, negative charged and large residues while less Ala, Ser, neutral, tiny and amide residues in the TPs. However, we only detected significant less Cys, Gln, and sulfur-containing and amide residues in the buried and partial buried area. This indicates more significant amino acids between TPs and MPs come from the external surface of proteins. These results were in good accordance with many previous reports that the differences were the most notable in protein surface, whereas not distinctive in the interior parts [4, 5, 7, 24, 28]. As for the secondary structure, we found the most obvious differences in the β-sheet. There were significant less Cys, Gln, Ser, tiny, sulfur-containing and amide residues in β-sheets. Meanwhile, we also observed significant more Glu, less Gln, tiny, amide residues in α-helix and only significant more Glu, and less amide residues in the coil. Most of our results were consistent with previous researches [8, 24, 28], although they were not as detailed as ours. From above-mentioned results, it is reasonable to believe the reliability of our definition on the significant amino acids (or amino acid groups) between TPs and MPs. Therefore, we can adopt it to identify the significant amino acids (or amino acid groups) between TPs and MPs according to their interaction degrees.

**Table 1.**
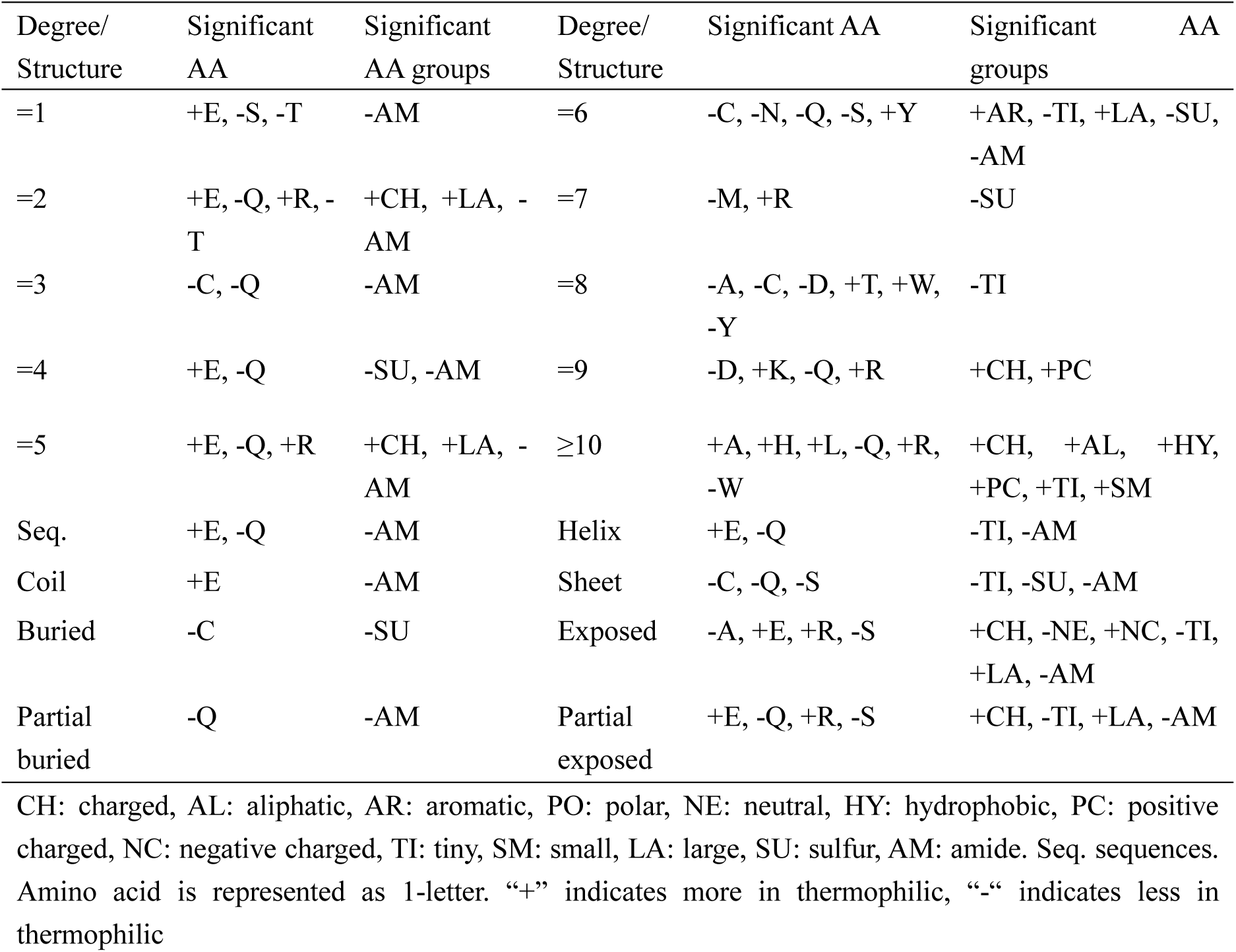
The significant amino acids and amino acid groups at various interaction degrees and structures.

We also listed the significant amino acids and amino acid groups at various interaction degrees in table 1. As can be seen that we detected more significant amino acid and amino acid groups when compared with the sequence. Among them, the significant residues and amino acid groups at degree 3 and 4 are nearly the same as sequence. As mentioned above, the Chi-squared distances between sequence and degree 3 and 4 are the minimum (Fig. 2). Taking these results together, we may conclude the amino acids differences in sequence between TPs and MPs mostly represented those who have three or four interactions with other residues.

For the significant amino acids, six of the 20 residues were assigned as significant (account 30% of the total) when the interaction degrees were eight and ten. Meanwhile, six of the 13 amino acid groups (with a proportion of 46.2%) also showed significant differences between TPs and MPs at the degree of 10 or above. As we strictly selected the homologous TPs and MPs, and only observed notable more Glu and less Gln in the sequences of TPs, these two residues also appeared in most of the interaction degrees as shown in table 1. In contrast, those significant amino acids detected at some interaction degrees could not find in the sequences. For example, less Ser and Thr in TPs at the interaction degree of 1; less Thr while more Arg in TPs at 2; and more Lys, Arg and less Asp in TPs at 9, respectively. Some previous works also identified these residues by comparing the sequences of TPs and MPs [8, 15, 24, 28]. These results indicated the method based on the interaction degree was more sensitive and could provide precise fingerprints about the significant amino acids (or groups). What needs to point out is that some amino acids play opposite roles at different interaction degrees. For instances, more Tyr in TPs at the interaction degree of 6 while less Tyr at 8; more Ala in TPs at 10 while less at 8; and more Thr in TPs while less at 1 and 2. These residues should be the potential mutation points when engineering the proteins for more stable at high temperature.

We also calculated the differences of amino acids between TPs and MPs according to the various ranges of interaction degrees and showed them in figure 3. We identified seven significant amino acids between TPs and MPs at various ranges of interaction degrees. Among them, two residues came from the low interaction degree, one from the medium degree and four from the high degree. At the low interaction degree, obvious more Glu and less Gln existed in TPs, which is the same as detected in the sequences. At the medium interaction degree, Cys was significantly lower in TPs, which has also reported before [15]. Interestingly, most of the significant amino acids were concentrated at the high interaction degree. We detected significantly more Phe, Lys, Leu and less Gln in TPs. More Lys, Leu and less Gln in TPs were commonly known [1, 8, 15] although most of them were not detected in the sequence. However, more Phe in TPs at high interaction degree borne out here is different. Although aromatic residues contribute positively to the conformational stability of TPs, only Tyr was frequently reported to have higher content in TPs [8, 15, 23, 24, 29]. As a contrast, rare reports about Phe contributing positively to the thermostability of proteins before. As we know, Phe is an aromatic residue, it can involve in the π-π stacking and π-cation in the aromatic clusters [30]. Moreover, the interaction between Phe and Cys/Met side chain at positions (*i*, *i* ± 4) can stabilize helices through interaction between aromatic electrons and sulfur atoms [31]. As thermophilic proteins usually have shorter sequence length [32] while more interactions [33], it is reasonable to expect the amino acids (such as Phe) involving in many interactions would be favorable in TPs. Some experimental results also confirmed the important role of Phe in the thermostability of proteins. For example, the T4 lysozyme mutant Ser 117 → Phe was isolated fortuitously and found to be more thermostable than wild-type by 1.1–1.4 kcal/mol [34]. While, the perturbation of the central residue (Phe 251) reduced the stability of the native structure of β-glucosidase [35].

**Fig. 3.**
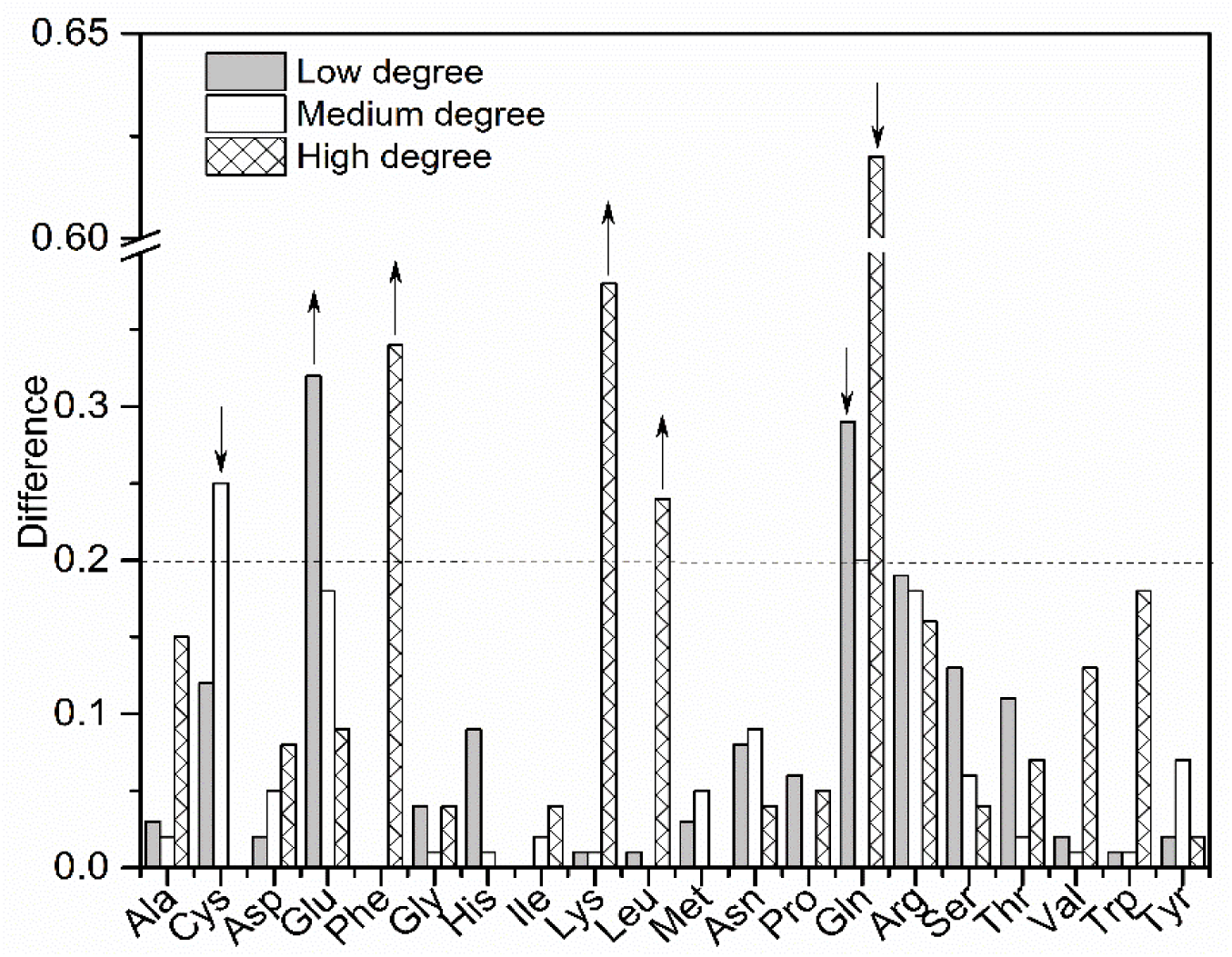
The differences of amino acids between thermophilic and mesophilic at various residue interaction degrees. The upwards arrow means more in TPs, and the downwards arrow means more in MPs.

We also calculated the differences of amino acids groups between TPs and MPs and showed them in figure 4. From it, we detected eight significant groups altogether, two groups came from the low interaction degree, one from the medium and five from the high. About 62.5% of the significant amino acid groups gathered at high interaction degrees. They were charged, positive charged, aliphatic, aromatic and small residues. It is well known these amino acid groups correlated positively to the protein thermostability [1, 2, 4, 8, 15, 23 However, we could not detect them in sequences with the same dataset. As mentioned earlier, the Chi-squared distances between sequence and the low interaction degree were the minimum (Fig. 2). Combining the two results, we can deduce the differences of amino acids in sequences between TPs and MPs could only represent those involved in low interaction degree and could not represent those high. Unfortunately, the amino acids involved in high interaction degrees are usually crucial to proteins. Thus, focus on the differences of the amino acids with high interaction degrees between TPs and MPs is of more importance and biological significance.

**Fig. 4.**
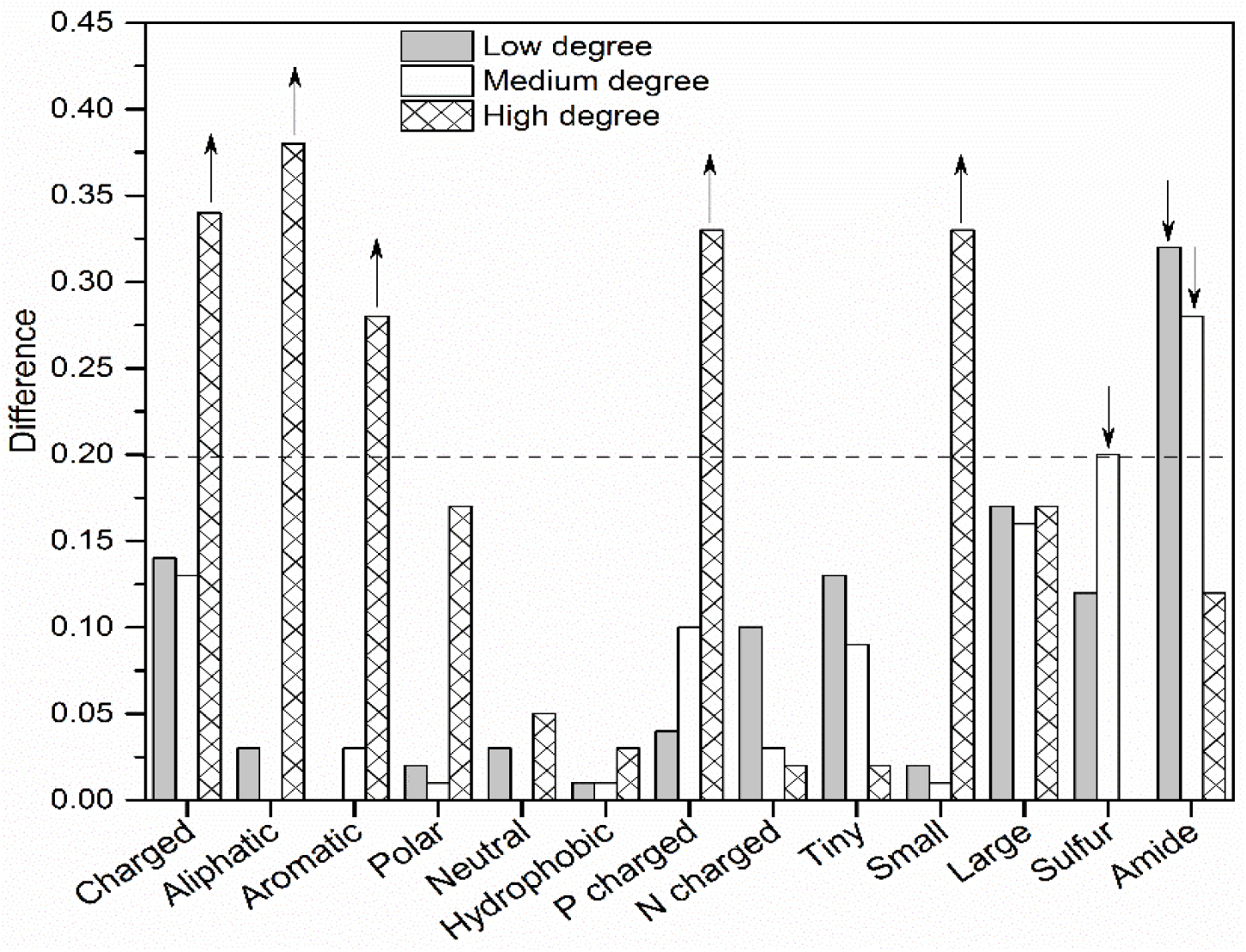
The differences of amino acid groups between TPs and MPs at various residue interaction degrees. The upwards arrow means more in TPs, and the downwards arrow means more in MPs.

Finally, as mentioned above, there were about 6.4% and 8.4% of the total amino acid in TPs and MPs did not involve in the interaction with other residues in our datasets. These non-involved residues may become the noise data when researchers tried to detect differences in sequences between TPs and MPs. Thus, this sometimes will lead to the conflict of their results. For example, some researcher observed more Gly in TPs while others found less Gly in TPs [27, 29]. Maybe because Gly is the most abundant residue among those who did not interact with other residues. As we found Gly accounted for 17.2% and 15.0% of the total residues in TPs and MPs that did not involve in the interaction with other residues. Besides, the interaction degree based method could remove the noisy amino acids in the sequence; it could be an important auxiliary tool for extracting features for predicting protein attributes [36], such as subcellular localization prediction, protein structural class, and membrane protein prediction and so on.

In short, detecting the significant amino acids between TPs and MPs based on their corresponding interaction degrees not only removed the noisy data in the sequences but also found more residues crucial for the thermostability of proteins. It could also distinguish the contribution of specific amino acid with various interaction degrees to the thermal stability of the protein. We will discuss this in the next paragraph.

### Differences of the interaction networks between the thermophilic and mesophilic aspartate transcarbamylase

To explain how a residue with various interaction degrees contributes differently to the thermostability of proteins according to their interaction network, we took thermophilic and mesophilic aspartate transcarbamylase as an example. The thermophilic one (PDB id: 1ML4) came from *Pyrococcus abyssi,* with an optimal growth temperature (OGT) of 97°C [16]. The mesophilic one (PDB id: 1Q95) came from *Escherichia coli*, with an OGT of 37°C [37]. The reason we chose them was that they shared 52.3% sequence identity and were close in sequence length (308 *vs* 310). The interactions (edges) within the molecule were 602 and 596 for 1ML4 and 1Q95 as shown in figure 5 A and B (drawn by Cytoscape [38]). The topology of the network was a little bit different. 1ML4 (TPs) has more compacted interaction in some area, while the interaction within 1Q95 was sparser and no obvious dense area detected (Fig. 5 A and B). Moreover, they also shared analogous structure according to the global alignment based on the interaction network as shown in figure 5 C and D. On the other hand, there were about 293 and 290 residues involved in the interaction, accounting 95.1% and 93.5% of the total residues (as shown in table 2). About 4.9% and 6.9% of the total residues did not involve in the interaction (the ratio is 6.4% and 8.4% for TPs and MPs in the whole dataset). This suggested more residues in MPs did not interact with others. These non-involved residues could be the potential mutations to increase the interactions, thus improving its thermostability. The residue content at low interaction degree is near, while the 1Q95 (mesophilic) have more residues with medium degree. Nevertheless, the differences of residue content at the high interaction degree are significant with *D*_*i*_ = 0.78. Eight residues with high interaction degree exist in 1ML4. They include Met-66, Tyr-226, Arg-297, Arg-107, Tyr-144, Arg-168, Met-265 and Met-298. However, only three in 1Q95, including Arg-105, Tyr-226 and Gln-288.

**Table 2.** Comparison of 1ML4 and 1Q95 according to the interaction degrees.

| **PDB ID** | **length** | **AAs with interactions** | **Low degree AAs** | **Medium degree AAs** | **High degree AAs** |
| --- | --- | --- | --- | --- | --- |
| 1ML4 (T) | 308 | 293 (95.1%) | 178 (60.8%) | 107(36.5%) | 8(2.7%) |
| 1Q95 (M) | 310 | 290 (93.5%) | 173 (59.7%) | 114(39.3%) | 3(1.0%) |

**Fig. 5.**
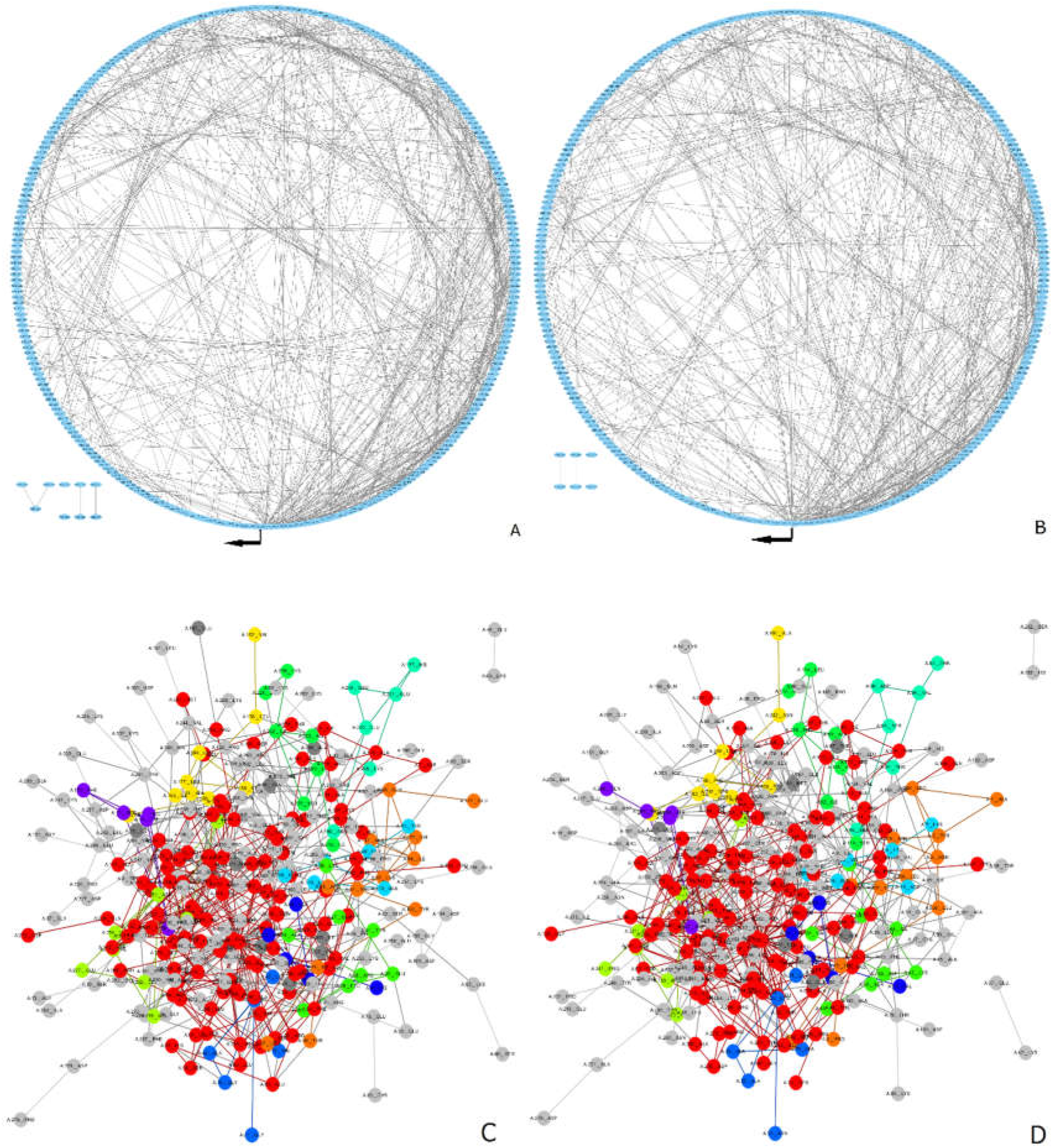
The residual interaction network of 1ML4 and 1Q95. A, B: Circular diagram representation of 1ML4 and 1Q95 interaction network; residues are presented as blue dots evenly distributed around the circle in a clockwise order starting from the black bar, which indicates the first residue involved in interaction with degree 1. Interaction between residues are represented by black lines. C, D: the global alignment of 1ML4 and 1Q95 based on residue interaction network. The top 10 largest common collected subgraphs (CCSs) are colored differently.

We adopted the MCODE [39] to find the highly interconnected regions that the residues with high interaction degrees involved and showed them in figure 6. We found five highly interconnected regions in 1ML4 while only two in 1Q95. This is due to more residues with high interaction degree in 1ML4. Besides, more residues with low or medium interaction degrees participated in these regions, leading more complex and concentrated interaction networks in 1ML4. Figure 6A showed the sub-network with the highest score (4.542) in 1ML4, there were 109 interactions (edges) and 49 residues (nodes) in the network, including the Tyr-226, Arg-297, Arg-107 and Arg-168 with high interaction degrees. Figure 6B showed the network with the second high score (4.0), there were 22 interactions and 12 residues in it, including the Met-66, Arg-297 and Met-298 with high interaction degrees. In the five sub-network of 1ML4, Arg-297 appeared 3 times, which was the most frequent residue. Undoubtedly, this Arg at position 297, which has the interaction degree of 10, plays a critical role in the thermostability of 1ML4. According to the annotation in PDB, Arg-297 located at the opposite of the binding pocket bottom, while Arg-107 and Arg-168 were part of the binding pocket. As mentioned in the introduction part, there were two argines (Arg-12 and Arg-152) not interacting with other residues in 1ML4, accounting 9.5% of the total argines. Their contributions to the thermostability of 1ML4 are undoubtly different from Arg-107, Arg-168 and Arg-297, which have the interaction degrees of 9, 9 and 10, respectively. In contrast, figure 6F showed the sub-network with the highest score (3.0) in 1Q95.

**Fig. 6.**
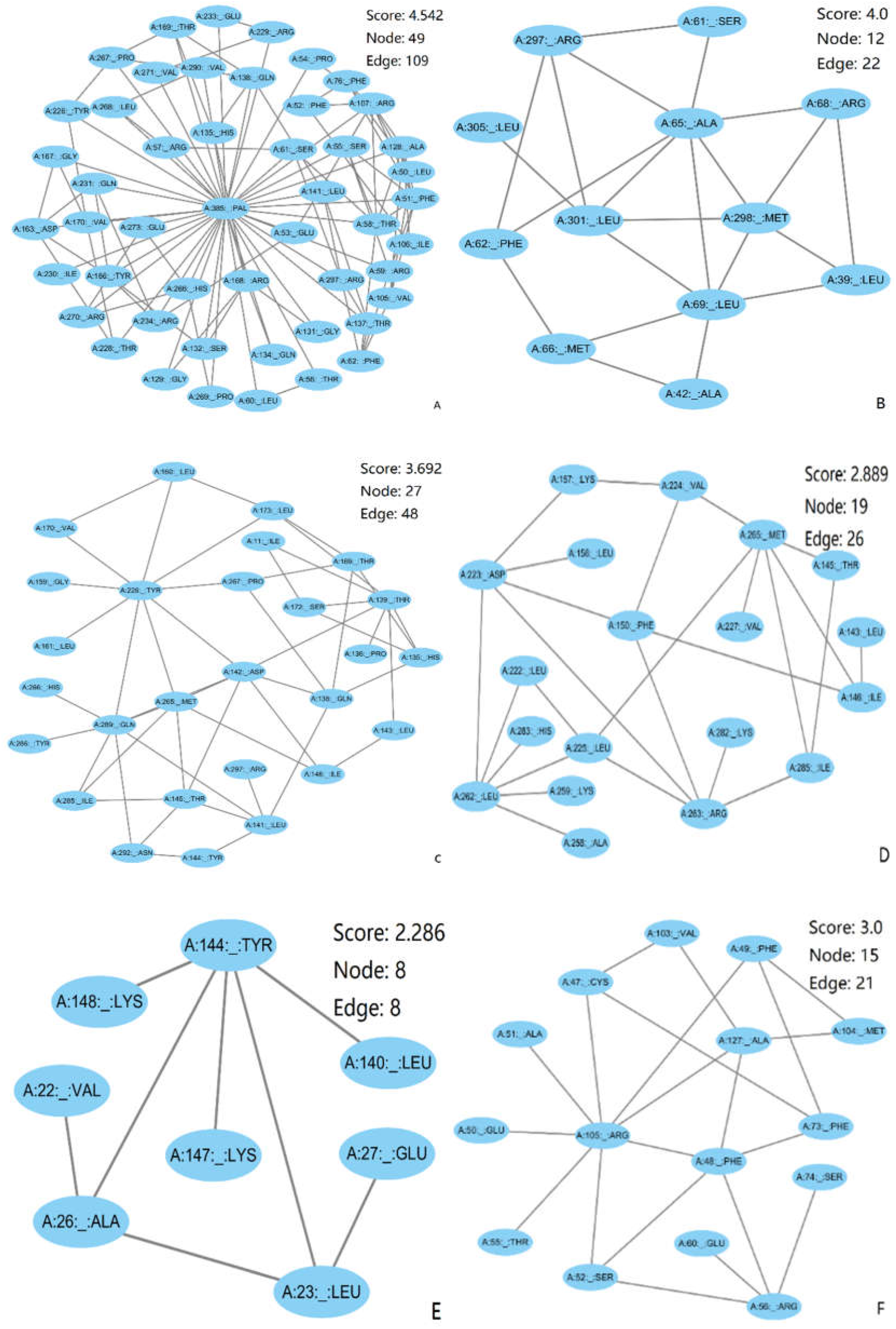
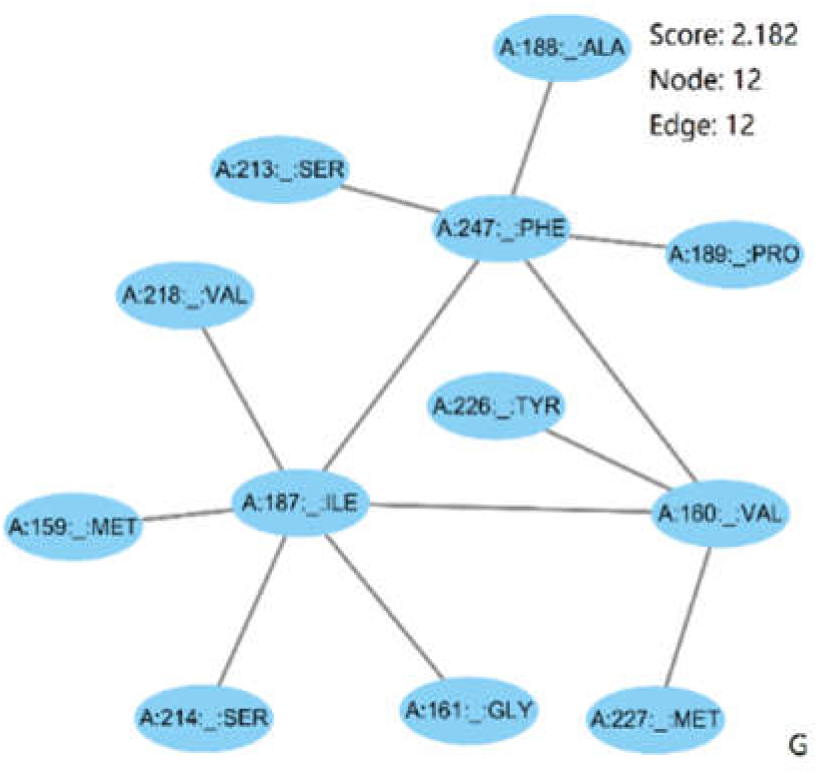
The highly interconnected regions the residues with high degrees (≥9) involved in 1ML4 and 1Q95. A-E: regions in 1ML4. The high degree residues involved include Met-66, Tyr-226, Arg-297, Arg-107, Tyr-144, Arg-168, Met-265 and Met-298. F-G: regions in 1Q95. The high degree residues involved include Arg-105, Tyr-226 and Gln-288.

There were only 21 interactions and 15 residues in the network, including the Arg-105 with high interaction degrees. Unlike 1ML4, all the argines in 1Q95 take part in the interaction with other residues, of which 10 argines come from low interaction degree, 4 from medium and 1 from high. Obviously, these argines contribute differently to the stability of 1Q95. Based on these facts, we firmly believe, the result will be more accurate and reliable if we remove the non-involved argines and compared the rest argines according to their interaction degrees. So do as other residues. Thus, we can detect more amino acids that are important when pursuing the influence of amino acids on the thermostability of proteins.

Moreover, figure S1 compares the interaction degree and evolutionary conservation score [40] of each residue in 1ML4 and 1Q95. There is a negative correlation between interaction degree and conservation score (r=0.42 and 0.43, respectively), meaning the amino acid with higher interaction degree remains more conserved. Generally, the conserved residues are less flexible and they are usually in the buried or surface region, whereas the disorder residues are the least conserved [41]. This has been proved by the results that both the hydrophobic core packing and the external residues were key factors for the stability of TPs [1, 2, 4, 7]. Meanwhile, most of the residues with high interaction degree are highly conserved. Their relative values of absolute surface areas (ASA) [42] were also the least, although some residues with low or medium degrees also have the least ASA (as shown in figure S2). About 81.8% of the non-involved amino acids (with the interaction degree of 0) have the ASA above 0.5, indicating these residues are mainly located on the surface of the aspartate transcarbamylase. They should be the first choice of mutation to improve the thermostability of this enzyme.

## Conclusion

In summary, we strictly selected 131 pairs of TPs and MPs, and detected the amino acids that showed significant differences based on their interaction degrees. We found more amino acids in the sequences of MPs did not interact with others; they might be the noisy data when identify the significant amino acids from sequences. Thus leading to obtain different sets of scarce and abundant amino acids for different research. For those residues involved in the interactions, the more interaction degrees of the residue, the more conserved of the amino acid, and the more important of it in maintaining the thermostability of proteins. We could observe much more significant amino acids (or amino acid groups) with high interaction degrees than in the sequences, indicating the interaction degree based method was more sensitive. We found significantly more Phe in TPs at high interaction degrees in the condition of no significant differences of Phe content existing in the sequences with the same dataset. As the interaction degree-based method can remove the noise of the sequence, it may act as an alternative tool to extract effective features to predict protein attributes in bioinformatics. Meanwhile, the residues with exact interaction degrees are more specific, which can provide more accurate and guiding information in engineering more thermostable proteins.

## Declarations

### Ethics approval and consent to participate

Not applicable

### Consent for publication

Not applicable

### Availability of data and material

All data generated or analyzed during this study are included in this published article [and its Additional file].

### Authors' contributions

Conceived and designed the experiments: HG YC GZ. Performed the experiments: HG YC. Analyzed the data: HG YC GZ. Wrote and revised the paper: HG and GZ.

## Acknowledgements

Not applicable

## Additional files

**Additional file 1** List of 131 pairs of thermophilic proteins and their mesophilic homologs

**Fig. S1** Comparison of interaction degree and the residue conservation score calculated with ConSurf (lower values indicate more conserved residues).

**Fig. S2** The correlation between ASA and interaction degree.

Table 1

CH: charged, AL: aliphatic, AR: aromatic, PO: polar, NE: neutral, HY: hydrophobic, PC: positive charged, NC: negative charged, TI: tiny, SM: small, LA: large, SU: sulfur, AM: amide. Seq. sequences. Amino acid is represented as 1-letter. “+” indicates more in thermophilic, “-” indicates less in thermophilic

